# Transcription Factor EB (TFEB) activity increases resistance of TNBC stem cells to metabolic stress

**DOI:** 10.1101/2023.05.30.542913

**Authors:** Milad Soleimani, Ria Goyal, Alexander Somma, Tamer S. Kaoud, Kevin N. Dalby, Jeanne Kowalski, S. Gail Eckhardt, Carla L. Van Den Berg

**Affiliations:** Interdisciplinary Life Sciences Graduate Programs, The University of Texas at Austin, Austin, Texas; Livestrong Cancer Institutes, Department of Oncology, Dell Medical School, The University of Texas at Austin, Austin, Texas; Division of Chemical Biology and Medicinal Chemistry, College of Pharmacy, The University of Texas at Austin, Austin, Texas; Division of Pharmacology and Toxicology, College of Pharmacy, The University of Texas at Austin, Austin, Texas

**Keywords:** TFEB, cancer stem cells, triple-negative breast cancer, UPR, autophagy

## Abstract

Breast Cancer Stem Cells (CSCs) are difficult to therapeutically target, but continued efforts are critical given their contribution to tumor heterogeneity and treatment resistance in Triple-Negative Breast Cancer (TNBC). CSC properties are influenced by metabolic stress, but specific mechanisms are lacking for effective drug intervention. Our previous work on TFEB suggested a key function in CSC metabolism. Indeed, TFEB knockdown (KD) inhibited mammosphere formation *in vitro* and tumor initiation/growth *in vivo*. These phenotypic effects were accompanied by a decline in CD44^high^/CD24^low^ cells. Glycolysis inhibitor 2-deoxy-D-glucose (2-DG) induced TFEB nuclear translocation, indicative of TFEB transcriptional activity. TFEB KD blunted, whereas TFEB (S142A) augmented 2-DG-driven UPR mediators, notably BiP/HSPA5 and CHOP. Like TFEB KD, silencing BiP/HSPA5 inhibited CSC self-renewal, suggesting that TFEB augments UPR-related survival. Further studies showed that TFEB KD attenuated 2-DG-directed autophagy, suggesting a mechanism whereby TFEB protects CSCs against 2-DG-induced stress. Our data indicate that TFEB modulates CSC metabolic stress response via autophagy and UPR. These findings reveal the novel role of TFEB in regulating CSCs during metabolic stress in TNBC.

**Financial Support:** This work was supported by CPRIT Grant RR160093 (to S.G. Eckhardt), CPRIT Grant RP210088 (to K.N. Dalby), UT College of Pharmacy Non-discretionary Funds (to C. Van Den Berg), and UT Graduate Continuing Fellowship (to M. Soleimani).

## INTRODUCTION

Triple-Negative Breast Cancer (TNBC) is a breast cancer subtype defined by the absence of Progesterone Receptor (PR), Estrogen Receptor (ER), and Human Epidermal growth factor Receptor 2 (HER2) (Won & Spruck, 2020). TNBC tumors are characterized by a high degree of heterogeneity, early metastatic onset, treatment resistance, and tumor relapse. Treatment of TNBC is more challenging than other breast cancer subtypes due to a lack of well-defined therapeutic targets and high tumor heterogeneity, leading to poor patient prognosis (Bai *et al*, 2021; Vagia *et al*, 2020)

Breast Cancer Stem Cells (CSCs) are a subpopulation of mostly quiescent cells that exhibit the capacity to self-renew and differentiate to reconstitute heterogeneous cell populations reflecting those of the original tumor. Mounting evidence points to TNBC harboring more CSCs than other breast cancer subtypes (Honeth *et al*, 2008; Ma *et al*, 2014). Indeed, CSCs contribute to the characteristic tumor heterogeneity, frequent metastasis, treatment resistance, and disease relapse observed with TNBC (Marra *et al*, 2020). This relatively small subpopulation is unique in terms of metabolic needs and plasticity, capable of adapting to metabolic and oxidative stress (Snyder *et al*, 2018). Understanding the metabolic mechanisms that govern CSC character and persistence could reveal new therapeutic avenues for TNBC.

The lysosomal membrane serves as a hub for metabolic signaling as lysosomes collect damaged organelles and recycle nutrients. Transcription Factor EB (TFEB) is a basic helix-loop-helix leucine zipper transcription factor best known for its key role in regulating lysosomal biogenesis and autophagy (Napolitano & Ballabio, 2016). Along with Melanocyte-Inducing Transcription Factor (MITF), TFEC, and TFE3, it belongs to the Microphthalmia Transcription factor E (MiT/TFE) family of transcription factors (Tan *et al*, 2022). TFEB phosphorylation, to a large extent, determines TFEB subcellular localization and activity. Once dephosphorylated, TFEB localizes to the nucleus and promotes the transcription of its target genes (Settembre *et al*, 2011). Aside from lysosomal biogenesis and autophagy, TFEB has been studied in the context of angiogenesis (Doronzo *et al*, 2019), immune response (Nabar & Kehrl, 2017), epithelial-mesenchymal transition (EMT) (Huan *et al*, 2005; Li *et al*, 2020), and cancer metabolism (Di Malta & Ballabio, 2017), among other processes in various disease-related models. Reprogrammed metabolism is a hallmark of cancer (Hanahan & Weinberg, 2011). Such alterations enable uncontrolled tumor growth or protect against oxidative and metabolic stress (DeBerardinis & Chandel, 2016).

TFEB lies at the intersection of various pathways directly associated with metabolic adaptation. mTORC1 regulates TFEB subcellular localization by phosphorylating S211, S142, and S122 (Vega-Rubin-de-Celis *et al*, 2017). Amino acid starvation inhibits mTORC1 activity, resulting in the nuclear localization of TFEB. Subsequently, TFEB activates the transcription of lysosomal biogenesis and autophagy genes which support nutrient catabolism (Martina *et al*, 2012). Cancers rely on glycolysis for energy production in a process called the Warburg effect. TFEB regulates multiple genes, including *HK1* (Hexokinase 1), *HK2*, *SLC2A1* (Solute Carrier family 2 member 1, a.k.a. GLUT1), and *SLC2A4* (a.k.a. GLUT4) that are involved in glucose metabolism (Mansueto *et al*, 2017). Glutamine is another source of metabolic support for tumors. It is directly involved in amino acid and nucleotide synthesis as a nitrogen donor and in cellular redox maintenance through glutaminolysis (Kim *et al*, 2021; Son *et al*, 2013). TFEB regulates glutamine metabolism by promoting the transcription of glutaminase (Kim *et al*., 2021). Thus, TFEB may be a nexus for cancer-associated metabolic stress and subsequent cell fate.

In this study, we report a novel role for TFEB in protecting triple-negative breast CSCs against metabolic stress. We have shown that glycolysis inhibitor 2-deoxy-D-glucose (2-DG) inhibits the CSC phenotype. 2-DG-induced stress triggers the unfolded protein response (UPR), increasing the expression of UPR-related genes such as BiP/HSPA5 and CHOP/DDIT3. This response is further augmented by nuclear TFEB. Additionally, TFEB protects CSCs against 2-DG-induced stress by promoting autophagy. Silencing BiP/HSPA5, a marker of breast CSCs (Conner *et al*, 2020), phenocopied TFEB knockdown in terms of mammosphere formation and CD24^low^/CD44^high^ enrichment, supporting that TFEB enriches CSCs by promoting UPR and autophagy.

## RESULTS

### TFEB promotes TNBC self-renewal *in vitro*

We have previously shown that knocking out TFEB reduces colony formation in TNBC cells (Soleimani *et al*, 2022). An initial analysis of TFEB expression across various breast cancer subtypes revealed that TNBC displays a higher expression of TFEB than normal mammary tissue and other breast cancer subtypes (Fig. 1A). In light of reports citing the ability of CSCs to endure metabolic and oxidative stress (Ciavardelli *et al*, 2014; Luo *et al*, 2018) and the function of TFEB in metabolic/oxidative response, we decided to examine the role of TFEB in self-renewal. We performed mammosphere formation and clonogenic assays on shRNA-mediated TFEB knockdown (KD) vs scramble cells. TNBC cell lines, HCC1806, HCC38, MDA-MB-231, and MDA-MB-157, were transduced with either scramble control or TFEB shRNA. Knocking down TFEB significantly inhibited secondary mammosphere formation in TNBC cell lines (Fig. 1B; Fig. EV1A & EV1B). This effect was consistent with the clonogenic assay results that also showed a dramatic decrease in colony formation in TFEB KD cells compared to the control (Fig. 1C; Fig. EV1C). It is widely documented that breast cancer tumors enriched in CD44^high^/CD24^low^ cells have a high tumor-initiating capacity (A & Lopes, 2017). Using data retrieved from Correlation AnalyzeR (Miller & Bishop, 2021), we observed that *TFEB* mRNA expression exhibited a direct correlation to *CD44* levels but no meaningful correlation to *CD24* (Fig. 1D). Moreover, knocking down TFEB depleted the CD44^high^/CD24^low^ population in human TNBC cell lines (Fig. 1E). The decline in mammosphere and colony formation and the concomitant decrease in CSC markers upon TFEB KD suggest that TFEB plays a role in breast cancer stem cell abundance.

**Figure 1.**
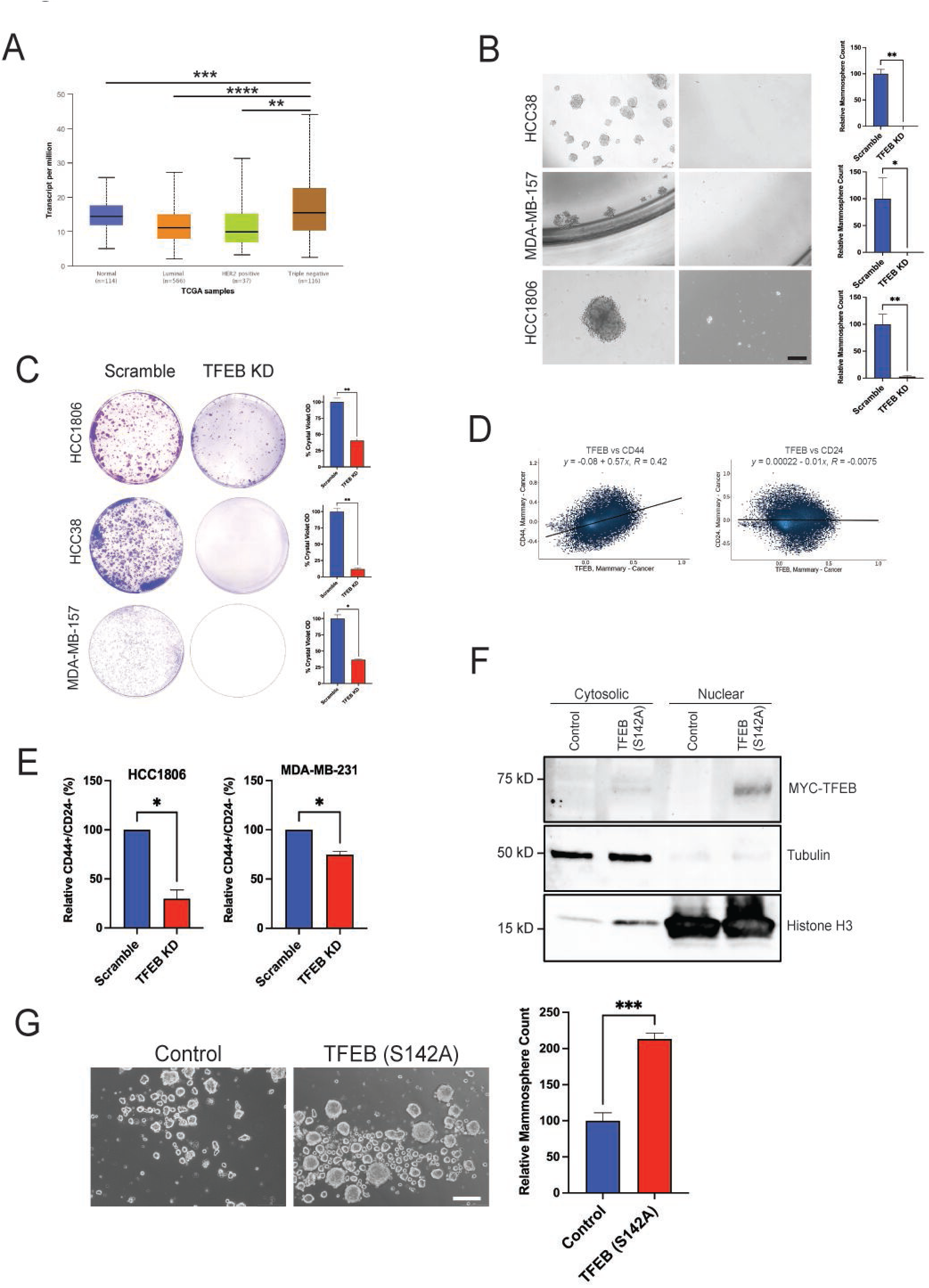
TFEB enriches TNBC CSC populations *in vitro.* **A)** UALCAN analysis of TFEB mRNA expression across breast cancer subtypes. (one-way ANOVA, Dunnett test) **B)** Mammosphere formation assay of indicated TNBC cell lines transduced with either scramble or TFEB shRNA and grown in mammosphere media for 7-10 days, followed by passaging to form secondary mammospheres. **C)** Clonogenic assay of indicated cell lines transduced with either scramble or TFEB shRNA. **D)** Scatter plots of TFEB vs. CD44 and TFEB vs. CD24 mRNA expression in breast cancer tumors as retrieved by Correlation AnalyzeR. **E)** Flow cytometric analysis of the CD44^high^/CD24^low^ fraction in indicated TNBC cell lines transduced with either scramble or TFEB shRNA. (Student’s *t-test*). **F)** Western blot analysis of cytosolic/nuclear fractions in HCC1806 expressing either empty pLenti-TrueORF or TFEB (S142A). **G)** Representative images and quantification of secondary mammospheres of HCC1806 cells expressing either empty control or TFEB (S142A). Scale bars: 200μm. *, *p* < 0.05; **, *p* < 0.01; ***, *p* < 0.001; ****, *p* < 0.0001

Having observed the impact of TFEB KD on cell self-renewal, we decided to assess the transcriptional effects of TFEB in cells. TFEB is phosphorylated by various kinases that regulate its subcellular localization and activity (Puertollano *et al*, 2018). Phosphorylation of TFEB by mTORC1 at S142 and/or S211 sequesters it in the cytoplasm, whereas the absence of phosphorylation at these sites results in the nuclear translocation of TFEB (Puertollano *et al*., 2018). As a transcription factor, nuclear TFEB induces the transcription of its target genes. To perform a gain-of-function study of active TFEB, we used a mutant TFEB construct harboring an alanine in place of serine at position 142 (Settembre *et al*., 2011). First, TFEB (S142A) was overexpressed in cells and validated by Western blot to have a predominantly nuclear localization (Fig. 1F). Additionally, ectopic expression of TFEB (S142A) increased mammosphere formation compared to control in TNBC cells (Fig. 1G).

### TFEB knockdown suppresses TNBC self-renewal *in vivo*

To assess the impact of TFEB KD on tumor growth kinetics *in vivo*, mice were orthotopically injected with either scramble control or TFEB KD HCC1806 cells. Over 20 days, TFEB KD cells displayed significantly slower growth than their scramble counterparts (Fig. 2A). We carried out a limiting dilution assay, the gold standard of cancer stem cell evaluation, to determine if TFEB KD affects tumor-initiating capacity *in vivo*. Mice were injected with either scramble control or TFEB KD HCC1806 cells serially diluted at 5×10^5^, 5×10^4^, 5×10^3^, or 5×10^2^. Tumor growth was monitored for 140 days following injection. ELDA (Extreme Limiting Dilution Analysis) analysis demonstrated that TFEB KD cells had an approximately 10-fold lower Tumor Initiating Cell (TIC) frequency compared to control cells (Fig. 2B). Hematoxylin-Eosin (H&E) staining revealed a pattern of cellularity in TFEB KD tumors that was less dense than that of control tumors (Fig. 2C).

**Figure 2.**
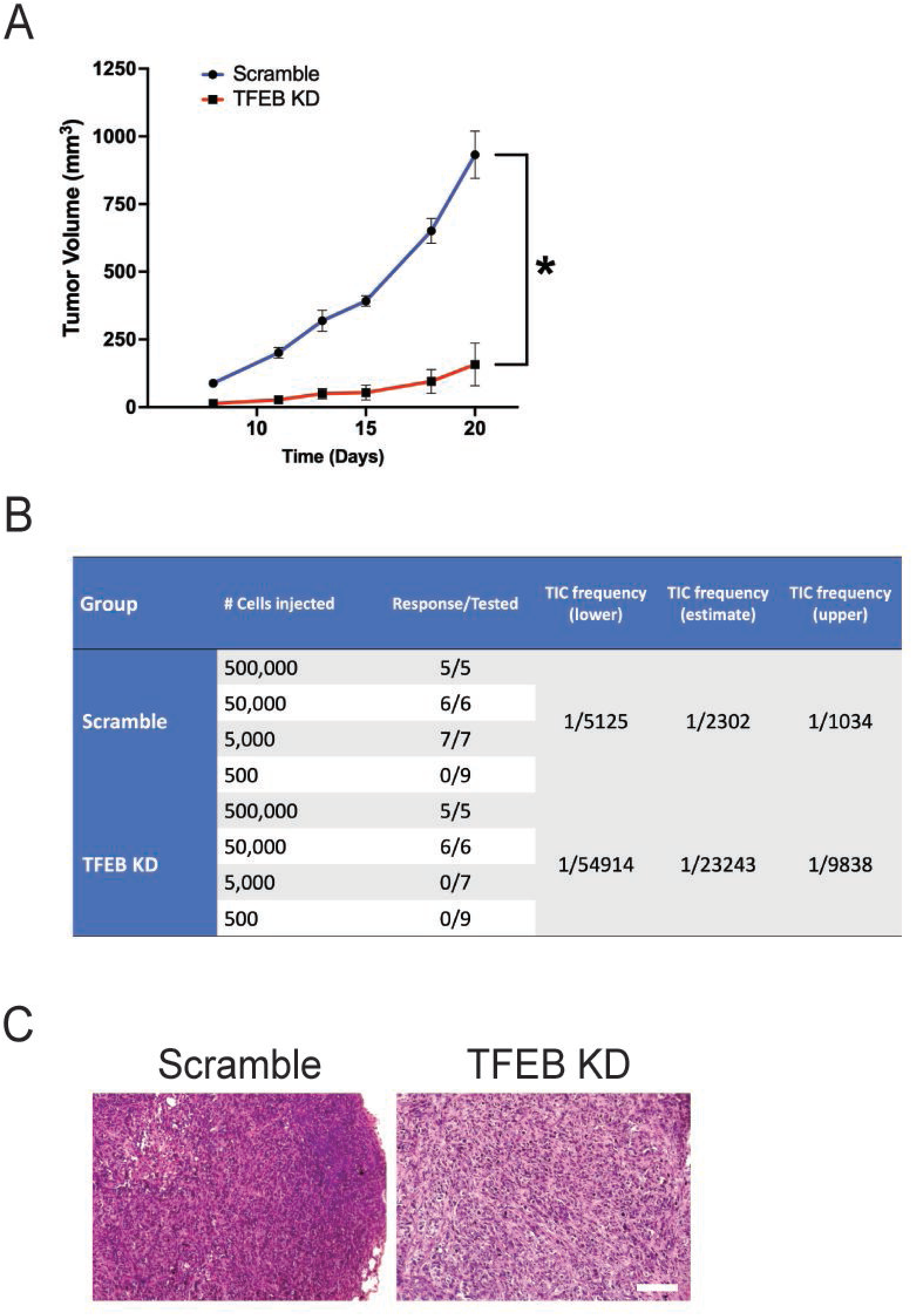
TFEB knockdown reduces TNBC tumor growth and self-renewal *in vivo*. **A)** Average tumor growth curves of HCC1806 cells transduced with either scramble (n=10) or TFEB shRNA (n=10) and injected orthotopically into nude mice. (Student’s *t-test*, *, *p* < 0.05) **B)** Tumor limiting dilution assay of HCC1806 cells transduced with scramble = or TFEB shRNA. NSG mice were orthotopically injected with 5✕10^5^ (n=5), 5✕10^4^ (n=6), 5✕10^3^ (n=7), or 5✕10^2^ (n=9) cells. **C)** Representative H&E images of either scramble control or TFEB KD HCC1806 xenograft tumors. Scale bar: 100μm.

### 2-DG inhibits TNBC growth and self-renewal

Normal cells depend primarily on oxidative phosphorylation for their metabolic needs, whereas cancer cells resort to aerobic glycolysis in a phenomenon called the Warburg effect (Liberti & Locasale, 2016). Evidence indicates that the transition from oxidative phosphorylation to glycolysis promotes cancer stemness in breast cancer (Dong *et al*, 2013). Others have demonstrated that blocking glycolysis with 2-DG reduces breast CSC populations (Luo *et al*., 2018). The mechanism whereby 2-DG impacts CSC character has yet to be determined. To assess the effect of 2-DG on cell viability, TNBC cell lines HCC1806, MDA-MB-231, MDA-MB-157, MDA-MB-453, HCC1937, HCC1395, HCC38, BT549, SW527, and HCC70 were treated with either vehicle or 0.3-20mmol/L 2-DG for 72h and analyzed using the CellTiter-Glo cell viability assay. All cell lines displayed a dose-dependent decline in cell viability in response to 2-DG (Fig. 3A). That being said, some cell lines, such as MDA-MB-157 and HCC70, were less sensitive than others, such as HCC1806 and HCC1937 (Fig. 3A). Similar trends were observed in colony formation where cells were treated with either vehicle or 1, 2, 5, 10, or 15 mmol/L 2-DG for 72h and recovered for 7-10 days (Fig. 3B; Fig. EV2B). To determine how 2-DG treatment impacts self-renewal *in vitro*, TNBC cell lines HCC1806, HCC38, and MDA-MB-157 were incubated in mammosphere media and treated with either vehicle or 2-DG. Each cell line was treated with its corresponding IC-50 dose of 2-DG derived from data in Fig. 3A. As expected, 2-DG suppressed self-renewal in every cell line tested, as illustrated by the significantly reduced mammosphere growth (Fig. 3C).

**Figure 3.**
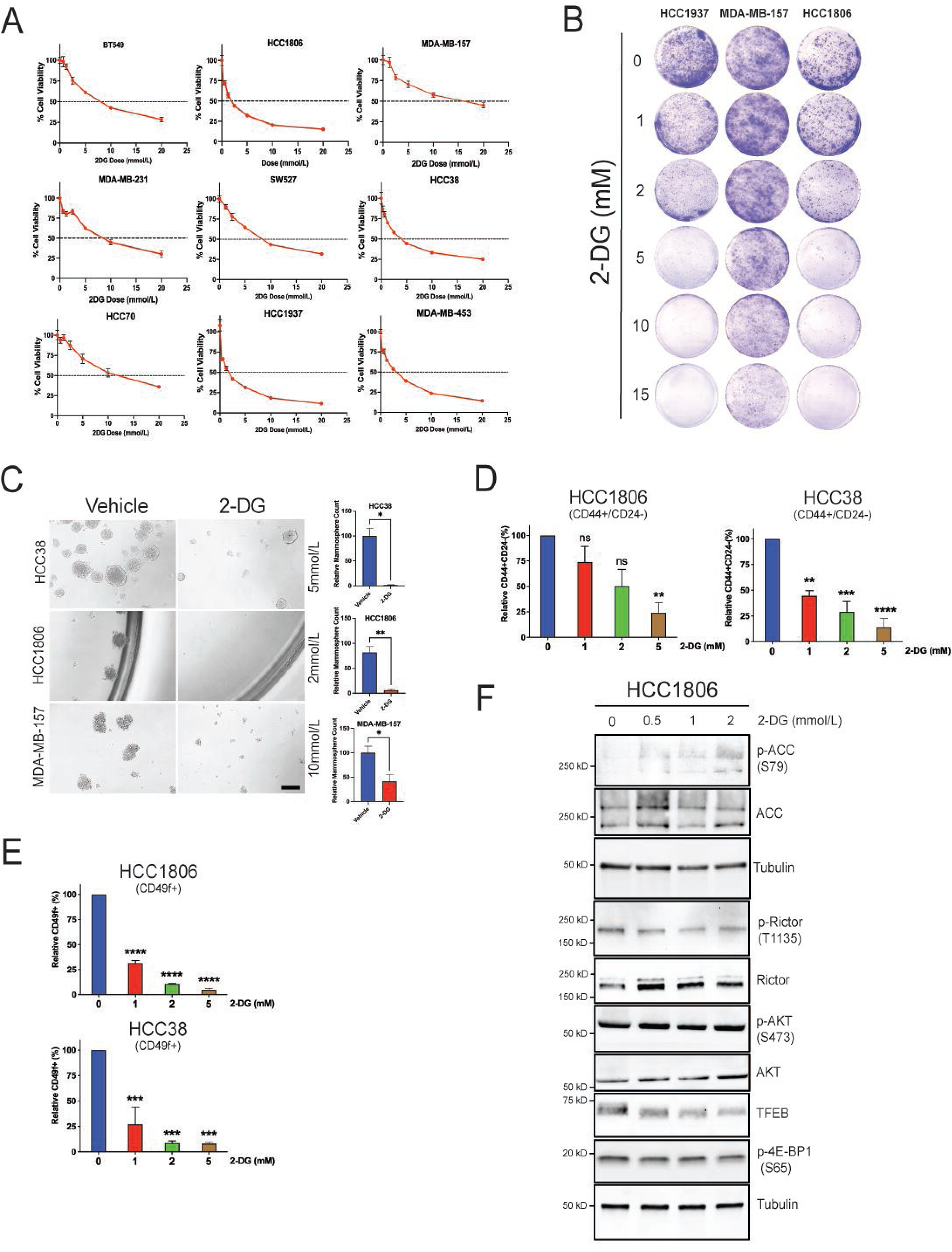
2-DG suppresses cell viability, and cancer stem cell phenotype in TNBC. **A)** Cell viability assay of indicated TNBC cell lines treated with either vehicle or 2-DG (0.3-20mmol/L) for 72h. **B)** Clonogenic assay of indicated TNBC cell lines treated with either vehicle or 2-DG (1, 2, 5, 10 & 20mmol/L) for 72h and recovered for 7-10 days. **C)** Mammosphere formation assay of indicated TNBC cell lines treated with either vehicle or 2-DG for 7-10 days. Scale bar: 200μm. **D)** Flow cytometric analysis of CD44^high^/CD24^low^ and **E)** CD49f fractions in indicated TNBC cell lines treated with either vehicle or 2-DG for 24h. (one-way ANOVA, Dunnett test) **F**) Western blot analysis of TFEB, p-4E-BP1 (S65), p-4-EBP1 (T37/46), p-p70S6K (T389), p-Rictor (T1135), p-AKT (S473), and p-ACC (S79) in HCC1806 treated with either vehicle or 2-DG for 24h. *p* < 0.05; **, *p* < 0.01; ***, *p* < 0.001; ****, *p* < 0.0001

To further verify the inhibition of CSC character by 2-DG, we examined various CSC biomarkers. We started by analyzing CD44^high^/CD24^low^ cells in HCC1806, HCC38, and HCC1937. Cells were incubated either in the absence or presence of 1, 2, 5 mmol/L 2-DG for 24h. There was a dose-dependent decline in CD44^high^/CD24^low^ cells (Fig. 3D; Fig. EV2B). Next, we interrogated another CSC biomarker CD49f (a.k.a. integrin alpha-6 (ITGA6)) in response to 2-DG treatment. Others have documented a close association between self-renewal and CD49f expression in human breast cancer (To *et al*, 2010). Our observations pointed to a striking reduction in CD49f levels, similar to those of CD44^high^/CD24^low^, in HCC1806 and HCC38 cells exposed to 2-DG for 24h (Fig. 3E). Together, we have shown, both phenotypically and using CSC biomarkers, that 2-DG-induced metabolic stress diminishes TNBC stem cell populations.

### 2-DG inhibits TFEB phosphorylation and induces TFEB nuclear translocation

We highlighted earlier that TFEB promotes cancer stemness in TNBC (Fig. 1; Fig. 2). TFEB plays an instrumental role in responding to metabolic stress. (Martina *et al*, 2016). These observations led us to investigate if TFEB regulates cellular response to glycolysis inhibitor 2-DG in TNBC. Preliminary Western blot analysis of TNBC cells treated with increasing concentrations of 2-DG revealed a gradual decrease in TFEB levels and a downward electrophoretic shift in TFEB bands (Fig. 3F). This shift in TFEB mobility is indicative of reduced phosphorylation. We have previously shown that inhibiting TFEB phosphorylation results in its nuclear translocation (Soleimani *et al*., 2022). Indeed, subcellular fractionation of cells treated with either vehicle or 2-DG revealed TFEB nuclear localization in the presence of 2-DG (Fig. EV3A). To confirm the effect of 2-DG on TFEB as purely glycolytic, we decided to glucose-starve cells for 3h, 8h, or 24h. The results were similar to those of 2-DG treatment in terms of both lowered TFEB expression and phosphorylation change (Fig. EV3B). TFEB activity and localization are regulated by multiple kinases, including mTOR, ERK, and AKT (Puertollano *et al*., 2018). To evaluate the effect of 2-DG on mTOR signaling in TNBC, we looked at mTORC1 target 4E-BP1 and mTORC2 targets AKT and Rictor. Surprisingly, 2-DG did not have a consistent impact across all cell lines on mTOR signaling at the concentrations used (Fig. 3F; Fig. EV3C). Another kinase upstream of TFEB is 5’-AMP-activated protein kinase (AMPK) (El-Houjeiri *et al*, 2019). AMPK activity increases in response to reduced ATP:AMP ratios induced by glycolytic stress. Unlike mTOR, AMPK showed a robust response to 2-DG treatment through phosphorylation of its well-known target Acetyl-CoA Carboxylase (ACC) (Fig. 3F; Fig. EV3C). These data demonstrate that 2-DG-driven stress increases TFEB electrophoretic mobility and AMPK activity. These effects closely mirror TFEB nuclear localization.

### 2-DG induces unfolded protein response

Unfolded protein response (UPR) is a stress mechanism to reduce the accumulation of unfolded or misfolded proteins causing endoplasmic reticulum (ER) stress. 2-DG is known to cause ER stress (Yu & Kim, 2010). UPR occurs via three major ER membrane stress sensors PERK (Protein kinase R (PKR)-like Endoplasmic Reticulum Kinase; a.k.a. EIF2AK), IRE1α (Inositol-Requiring transmembrane kinase/Endoribonuclease 1α), and ATF6 (Activating Transcription Factor 6) (Hetz *et al*, 2020). Various components of UPR are closely associated with breast CSC character, suggesting that UPR preserves CSC populations. Inhibition of ATF6 and PERK suppresses mammosphere formation (Li *et al*, 2018). Knockdown of X-box Binding Protein 1 (*XBP1*) reduces CD44^high^/CD24^low^ enrichment in TNBC (Chen *et al*, 2014). Overexpression of Binding immunoglobulin Protein (BiP; a.k.a. GRP78, HSPA5) increases CD44^high^/CD24^low^ cells and upregulates CSC-associated genes (Conner *et al*., 2020). We measured the change in the expression of several UPR markers caused by 2-DG exposure, namely *XBP1*, DNA damage-inducible transcript 3 (*DDIT3*, a.k.a. CHOP), Activating Transcription Factor 4 (*ATF4*), Protein Disulfide Isomerase Family A member 2 (*PDIA2* a.k.a. PDI), ER Degradation Enhancing α-Mannosidase like protein 1 (*EDEM1*), Protein Phosphatase 1 Regulatory subunit 15A (*PPP1R15A*), Heat Shock 70 kDa Protein 5 (*HSPA5*), and Activating Transcription Factor 6 (*ATF6*) in HCC1806 and HCC38 using qRT-PCR (Fig. 4A). We further confirmed UPR activation by 2-DG using Western blot. To that end, HCC1806, HCC70, BT549, MDA-MB-157, and HCC1937 cells were treated with either vehicle or increasing concentrations of 2-DG. The results revealed UPR induction as shown by an upregulation of PERK, PDI, BiP, CHOP, and IRE-1α (Fig. 4B). Tunicamycin (TM), a well-documented ER stress inducer, mostly mirrored the effect of 2-DG on UPR (Fig. 4C).

**Figure 4.**
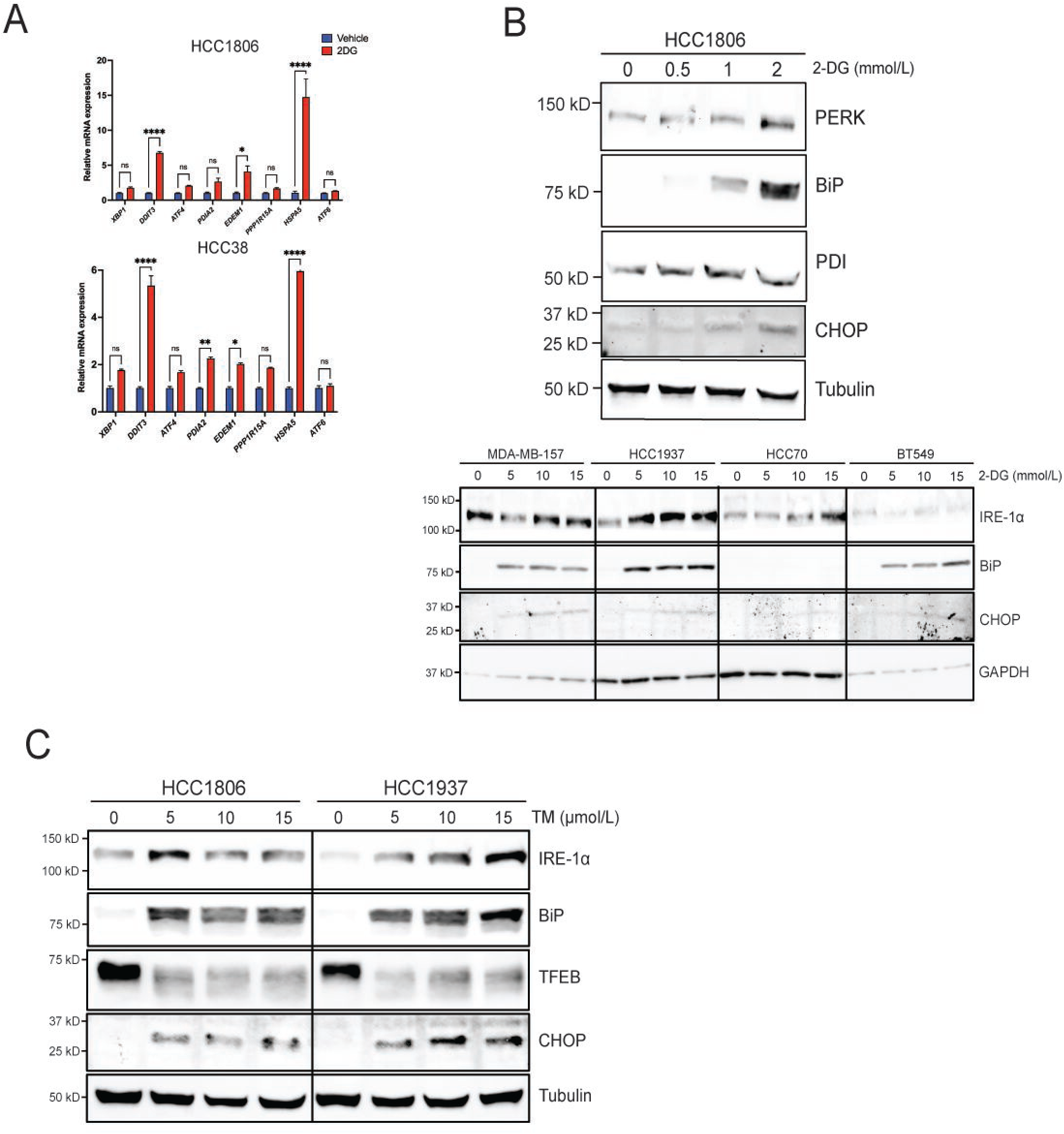
2-DG induces unfolded protein response (UPR). **A)** qRT-PCR analysis of indicated UPR markers in HCC1806 and HCC38 treated with either vehicle or 2-DG for 24h. (two-way ANOVA, Sidak test) **B)** Western blot analysis of UPR markers in indicated TNBC cell lines treated with either vehicle or 2-DG (2mmol/L) for 24h. **C)** Western blot analysis of UPR markers in indicated TNBC cell lines treated with either vehicle or increasing concentrations of tunicamycin. Veh: Vehicle; TM: Tunicamycin *, *p* < 0.05; **, *p* < 0.01; ***, *p* < 0.001; ****, *p* < 0.0001

### TFEB mediates 2-DG-driven UPR

First, we investigated the role of TFEB in responding to 2-DG-induced metabolic stress and the resulting impact on CSCs. Normally, TFEB resides in the cytoplasm bound to the lysosomal membrane through interaction with a component of the Ragulator complex but localizes to the nucleus under various types of stress. To determine the significance of TFEB subcellular localization, we generated cell lines expressing either predominantly nuclear or cytosolic TFEB: 1) TFEB (S142A) is a constitutively nuclear TFEB mutant (Fig. 1F) and 2) RagC (S75L) sequesters TFEB in the cytoplasm (Fig. 5A). First, we performed a mammosphere formation assay with cells transduced with TFEB (S142A), RagC (S75L), or empty vector treated with either vehicle or 2-DG and incubated for 7-10 days. Stable overexpression of TFEB (S142A) lowered 2-DG sensitivity in TNBC mammospheres compared to control. In contrast, overexpression of RagC (S75L) enhanced 2-DG cytotoxicity (Fig. 5B). These findings align with the observation that nuclear TFEB positively regulates CSC self-renewal in TNBC.

**Figure 5.**
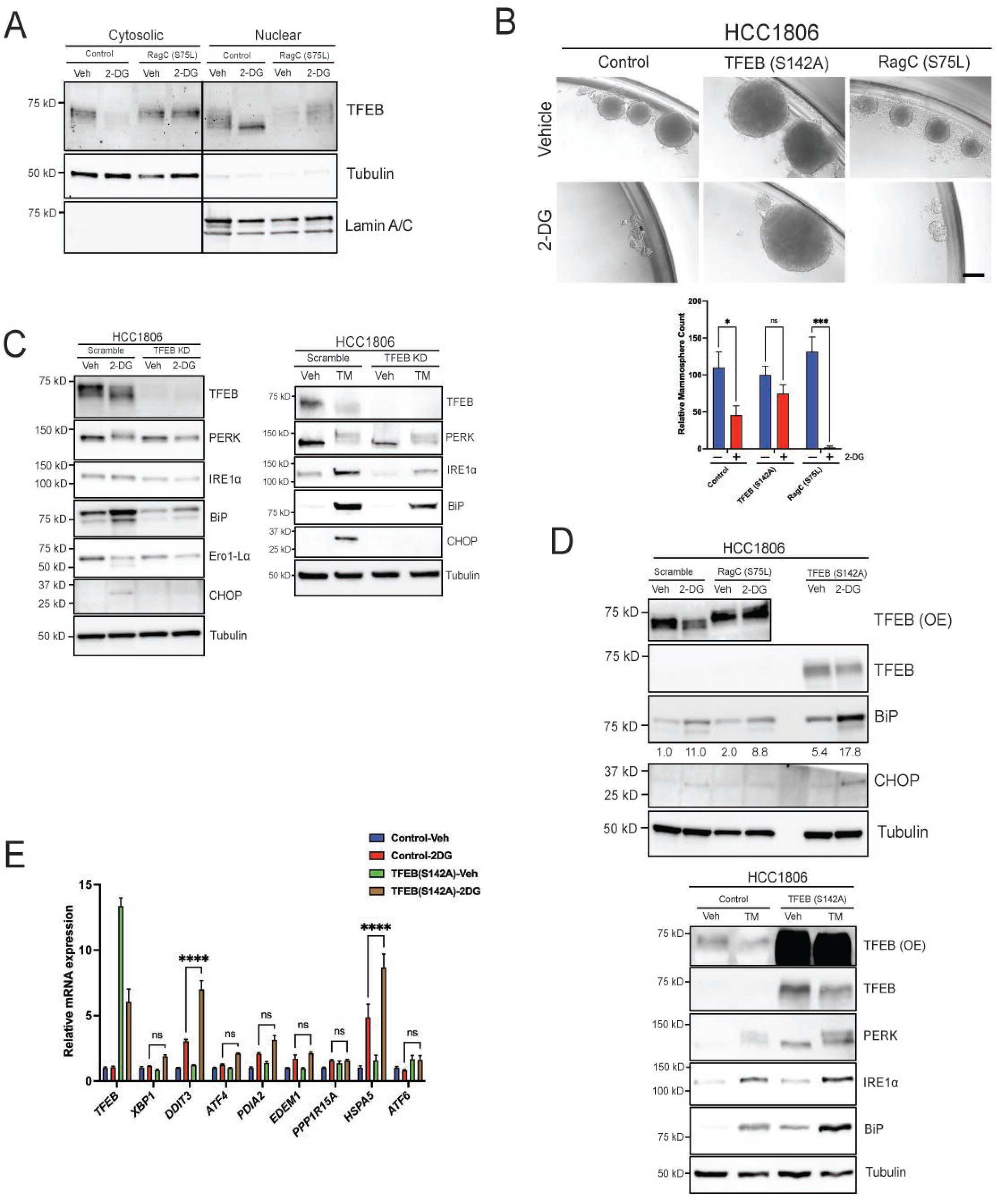
TFEB mediates 2-DG-driven UPR. **A)** Western blot analysis of nuclear/cytosolic fractions of either pLenti or RagC (S75L) treated with either vehicle or 2-DG for 24h. Tubulin was used a cytosolic marker and Lamin A/C as a nuclear marker. **B)** Mammosphere formation assay of empty pLenti, TFEB (S142A), or RagC (S75L) HCC1806 treated with either vehicle or 2-DG. (two-way ANOVA, Sidak test) Scale bar: 200μm. **C**) Western blot analysis of UPR markers in either scramble or TFEB KD and **D)** empty pLenti, RagC (S75L), or TFEB (S142A) treated with either vehicle, 2-DG, or tunicamycin for 24h. **E)** qRT-PCR analysis of UPR markers in either empty pLenti or TFEB (S142A) cells treated with either vehicle or 2-DG (2mmol/L) for 24h. Veh: Vehicle; TM: Tunicamycin *, *p* < 0.05; **, *p* < 0.01; ***, *p* < 0.001; ****, *p* < 0.0001

Given that both TFEB and UPR enhance cancer stemness (Liang *et al*, 2021; Spaan *et al*, 2019) and are responsive to metabolic stress, we hypothesized that TFEB modulates 2-DG-induced UPR. To test this hypothesis, we compared UPR activation by 2-DG in scramble control and TFEB KD cells. A Western blot analysis showed that silencing TFEB blunted 2-DG upregulation of UPR markers (Fig. 5C). Indeed, overexpression of constitutively nuclear TFEB (S142A) augmented upregulation of BiP and CHOP at both protein (Fig. 5D) and mRNA (Fig. 5E) levels. Interestingly, TFEB KD diminished UPR induction, and TFEB (S142A) enhanced it in cells treated with tunicamycin (Fig. 5C & 5D). As clinical correlates, *TFEB* mRNA expression in TNBC patient samples showed a direct correlation of TFEB expression to *EIF2AK*, *DDIT3*, and *HSPA5* (Fig. EV3D).

### BiP/*HSPA5* knockdown suppresses CSC phenotype

We have shown that 2-DG activates a TFEB-UPR axis while suppressing cancer stemness. Others have reported that BiP/*HSPA5* overexpression (a key UPR-associated gene) increases CD44^high^/CD24^low^ cells in breast cancer (Conner *et al*., 2020). We aimed to determine if silencing BiP/*HSPA5*, also a TFEB-responsive gene, impacts TNBC self-renewal. To validate the shRNA-mediated BiP/*HSPA5* KD, we performed a Western blot on scramble and BiP/*HSPA5* KD cells treated with either vehicle or 2-DG for 24h. BiP/*HSPA5* KD attenuated BiP/*HSPA5* and CHOP upregulation in response to 2-DG (Fig. 6A). In untreated TNBC cells, BiP/*HSPA5* KD inhibited self-renewal as indicated by significantly reduced colony and mammosphere formation (Fig. 6B & 6C). Further, an analysis of CSC biomarkers revealed a decline in CD44^high^/CD24^low^ cells upon BiP/*HSPA5* KD, consistent with a diminished CSC population (Fig 6D).

**Figure 6.**
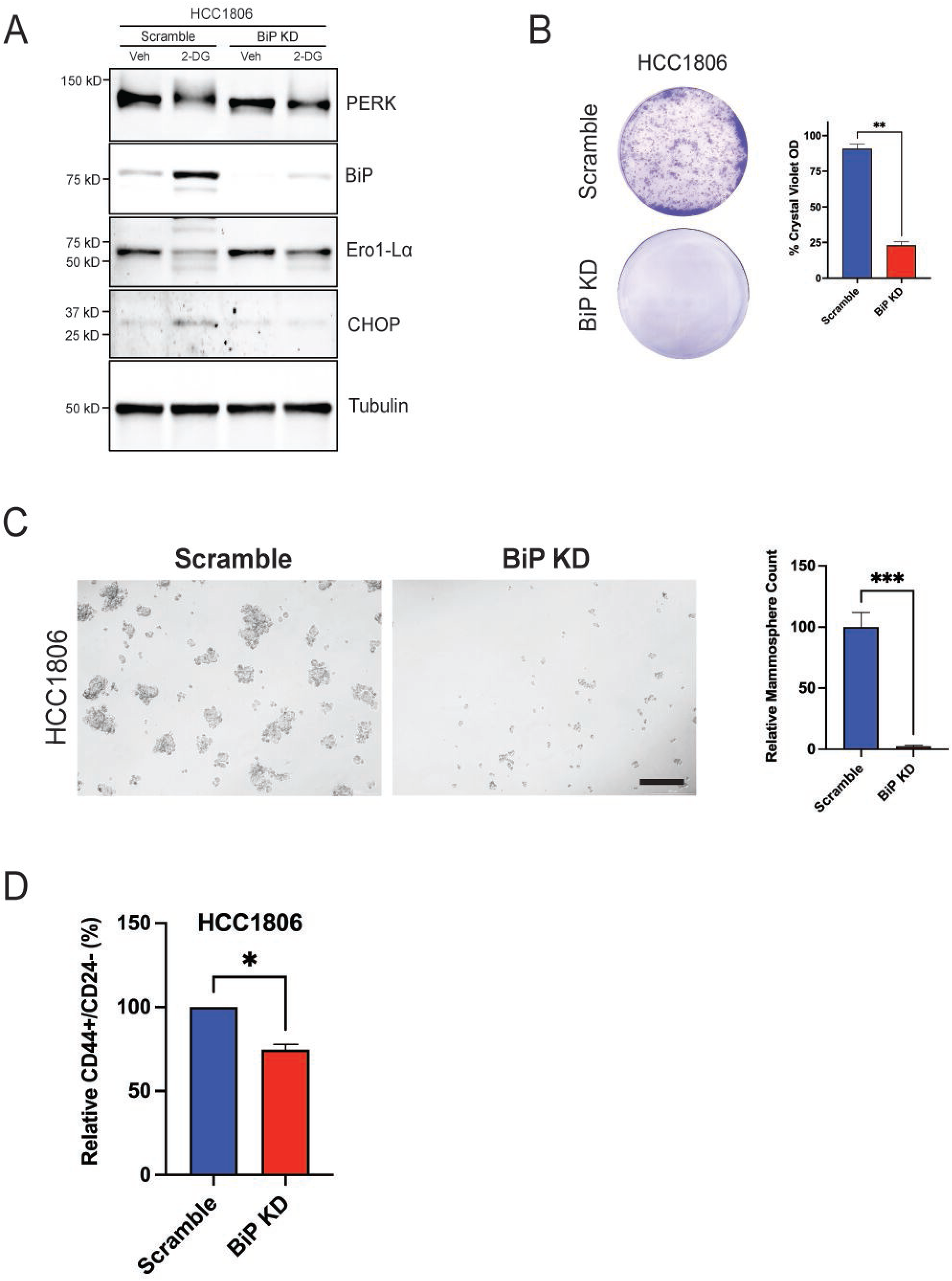
BiP/HSPA5 knockdown suppresses TNBC self-renewal. **A**) Western blot of indicated UPR markers in either scramble or BiP/HSPA5 KD cells treated with either vehicle or 2-DG for 24h. **B)** Clonogenic assay and **C)** mammosphere formation assay of indicated cell lines transduced with either scramble or BiP/HSPA5 shRNA. Scale bar: 200μm. **D**) Flow cytometric analysis of CD44^high^/CD24^low^ cells in HCC1806 transduced with either scramble or TFEB shRNA. (Student’s *t-test*, *, *p* < 0.05). Veh: Vehicle

### TFEB and UPR promote autophagy in response to 2-DG

Thus far, our data show that 2-DG induces UPR and TFEB activity. However, we lacked an explanation as to how TFEB rescues CSCs from 2-DG-induced stress. CSCs and tumor cells utilize UPR as a survival response and can engage autophagy (ER-phagy and mitophagy) to clear unfolded proteins and damaged organelles (Senft & Ronai, 2015). We anticipated TFEB might shift cells toward an autophagic survival response since TFEB directly regulates the expression of autophagy-and lysosome-related genes (Settembre *et al*., 2011). We tested if 2-DG induces autophagy, as shown by increased p62 and LC3-II compared to vehicle controls. Indeed, there was an increase in p62 and LC3-II levels upon 2-DG treatment. TFEB KD reduced the autophagic response to 2DG (Fig. 7A). Additionally, co-treatment with the PERK inhibitor ISRIB to reduce UPR showed a similar response to TFEB KD, attenuating the autophagic response to 2-DG (Fig. 7B). We validated the effect of ISRIB on autophagy by assessing LC3 using ICC (Fig. 7C). Hydroxychloroquine (HXQ) was used as a positive control to cause LC3 puncta. Together, these data support that 2-DG and TFEB increase the expression of several UPR genes while TFEB promotes autophagy to sustain cancer and CSC populations.

**Figure 7.**
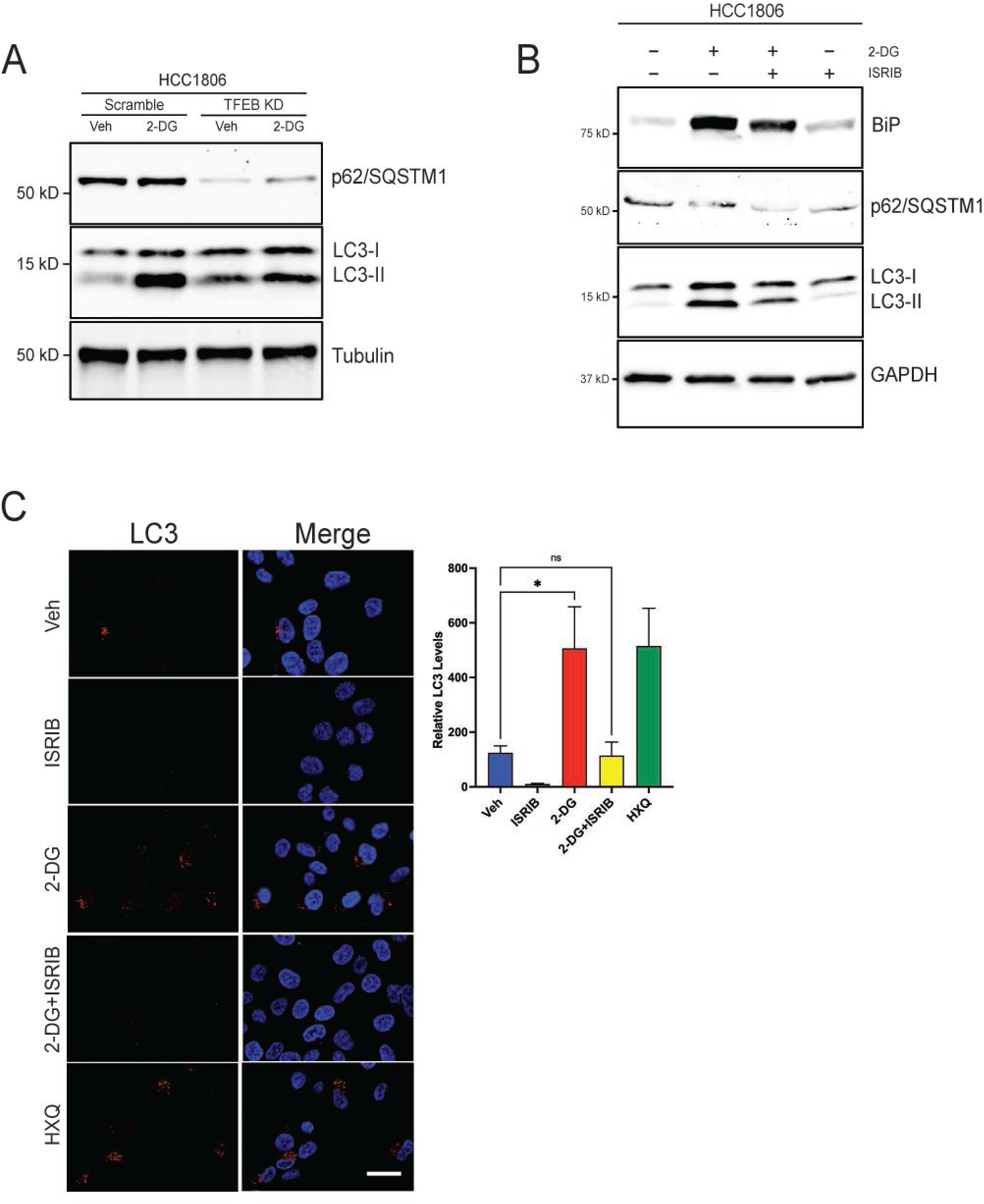
TFEB induces autophagy in response to 2-DG. **A**) Western blot of p62/SQSTM1 and LC3 in either scramble or TFEB KD treated with either vehicle or 2-DG for 24h. **B**) Western blot of LC3, p62/SQSTM1, and BiP in cells treated with vehicle, 2-DG, and/or ISRIB for 24h. **C**) Representative ICC images of LC3 and DAPI in HCC1806 cells treated with vehicle, 2-DG-ISRIB, 2-DG+ISRIB, or hydroxychloroquine. (one-way ANOVA Dunnett test). Scale bar: 25μm. Veh: Vehicle. HXQ: Hydroxychloroquine. *, *p* < 0.05; **, *p* < 0.01; ***, *p* < 0.001; ****, *p* < 0.0001

## DISCUSSION

The unique metabolic landscape of CSCs represents a therapeutic opportunity. Cancer cells undergo metabolic alterations to meet their energetic needs for growth. These alterations result in a glycolytic mode of metabolism accompanied by an increase in glucose uptake (Hanahan & Weinberg, 2011). CSCs, on the other hand, are metabolically flexible in response to the tumor microenvironment (Snyder *et al*., 2018). One hypothesis is that CSCs follow the hierarchical pattern of normal cells where differentiation is closely associated with a switch from glycolysis to oxidative phosphorylation (Sancho *et al*, 2016). Others have postulated that CSC character is driven by metabolic reprogramming in cells (Menendez & Alarcon, 2014). A comprehensive model of non-CSC-to-CSC transition showed that intermediary metabolites, such as α-ketoglutarate and Acetyl-CoA, govern the epigenetic regulation of genes involved in CSC character (Menendez & Alarcon, 2014). In breast cancer, researchers have documented the significance of glycolytic shift and enhanced macromolecule biosynthesis to cancer stemness maintenance (Dong *et al*., 2013). Several seemingly contradictory reports point to either glycolysis (Ciavardelli *et al*., 2014; Zhou *et al*, 2011) or oxidative phosphorylation (Janiszewska *et al*, 2012) as the preferred mode of CSC metabolism. Understanding how metabolic stress modulates CSC dynamics is key to developing an effective treatment that targets the entire CSC population.

Given the central role that autophagy and lysosomes play in cellular metabolism and that TFEB is a master regulator of genes involved in metabolic adaptation, we decided to explore whether TFEB controls CSC populations with or without metabolic stress. Herein, the knockdown of TFEB in TNBC cells strongly inhibited mammosphere formation and tumor initiation. First, we utilized CSC biomarkers CD44^high^/CD24^low^ as well as *in vitro* and *in vivo* functional assays to assess the role of TFEB in CSCs regardless of metabolic stress. We also found TFEB pivotal in safeguarding CSCs against metabolic stress. Our study specifically indicates that TFEB attenuates the suppressive effects of 2-DG-induced stress on TNBC self-renewal by promoting autophagy and UPR. Autophagy involves, among other things, the lysosomal metabolism of sulfur amino acids, which contributes to the cellular cysteine reservoir for antioxidant defense and adaptation (Matye *et al*, 2022). mTOR-driven aberrant suppression of autophagy sensitizes cisplatin-resistant lung cancer to 2-DG (Gremke *et al*, 2020). Another instance of TFEB responding to metabolic stress is its induction of mitophagy to counteract metabolic/mitochondrial stress-driven Ca^2+^ release to support pancreatic β-cell function (Park *et al*, 2022). Altogether, TFEB and the associated metabolic stress response machinery represent a potential vulnerability in CSCs and a promising area for therapeutic exploration. It is also important to point out that the RNA or protein abundance of TFEB is less likely to serve as a biomarker of its activity than its subcellular localization. Our studies and others bear out this conclusion (Zhu *et al*, 2021).

We have demonstrated that 2-DG suppresses mammosphere formation and CD44^high^/CD24^low^ cells associated with mesenchymal-like CSCs in TNBC. It bears mentioning that chronic metabolic stress promotes breast cancer stemness in a Wnt-dependent fashion (Lee *et al*, 2015), and that 2-DG promotes ALDH+ epithelial-like CSCs while inhibiting invasive mesenchymal-like CSCs (Luo *et al*., 2018). Hyperactivation of UPR and/or autophagy as a consequence of 2-DG treatment could lead to increased CSCs. Furthermore, it is critical to acknowledge that each CSC assay alone is insufficient to accurately assess CSC character. Therefore, we have used a combination of CD44^high^/CD24^low^ and CD49f+ cells as biomarkers. Additionally, we have utilized functional assays in the form of clonogenic and mammosphere assays *in vitro* and a tumor-limiting dilution assay *in vivo*. Using several different methods is vital to ensure the reliability and robustness of CSC data.

A mechanistic analysis of our results revealed that TFEB-directed 2-DG response occurs via UPR. In breast cancer, UPR is documented to stimulate cancer stemness (Liang *et al*., 2021). 2-DG induced UPR in TNBC cells in a dose-dependent fashion. This effect was mitigated upon TFEB KD and amplified upon TFEB (S142A) overexpression. We have shown that BiP/*HSPA5* and CHOP/*DDIT3* are the most consistently upregulated UPR markers at both mRNA and protein levels. Further, these were the two most altered markers due to either TFEB KD or overexpression. There are several UPR markers implicated in CSC regulation. BiP localized to the cell surface promotes CSC phenotype and metastasis in breast cancer (Conner *et al*., 2020). It is upregulated in bone marrow-derived disseminated breast tumor cells displaying cancer stem cell character (Bartkowiak *et al*, 2010). A meta-analysis of BiP and its clinicopathological potential in breast cancer found a correlation between high BiP expression and HER2 and basal-like subtypes as well as metastatic tumors (Direito *et al*, 2022). XBP1 is critical for the tumorigenicity, progression, and relapse of TNBC, where it forms a complex with HIF1α (Hypoxia Inducible Factor 1 subunit α) to maintain CSCs (Chen *et al*., 2014). TNBC/basal-like tumors have higher levels of TFEB than the other breast cancer subtypes. This renders studying TFEB and its regulation of metabolic stress and CSCs in TNBC more clinically relevant. Although TNBC patients generally respond to chemotherapy, they have an earlier relapse and a more frequent recurrence than the other breast cancer subtypes (Zagami & Carey, 2022). This is partly attributable to the dormant CSC population that is insensitive to most therapies but gives rise to tumor heterogeneity. Thus, new therapeutics must target promiscuous CSCs to overcome the barriers associated with tumor heterogeneity, treatment resistance, metastasis, and tumor recurrence.

In conclusion, we have uncovered a novel metabolic stress response mechanism where TFEB sustains CSCs by upregulating UPR and autophagy in TNBC. TFEB depletion suppressed self-renewal *in vitro* and *in vivo*, and overexpression of active TFEB thwarted 2-DG-induced mammosphere inhibition. Mechanistically, TFEB KD cells showed a diminished UPR response to 2-DG, while TFEB (S142A)-overexpressing cells had a more robust response than the corresponding controls. The key role of TFEB during metabolic stress appears to be as a UPR-responsive gene that promotes cell survival by enhancing autophagy and other key UPR-related genes that regulate TNBC CSCs, namely BiP. Our limited clinical analyses further support a metabolic gene signature involving TFEB regulation of CSC and UPR markers. This further supports pursuing these pathways to better understand CSC biology and potential new targets for treating TNBC.

## MATERIALS AND METHODS

### Cell culture and reagents

TNBC cell lines MDA-MB-231, MDA-MB-157, MDA-MB-453, HCC1806, HCC70, HCC38, SW527, HCC1395, HCC1937, and BT549 were obtained from the American Type Culture Collection (ATCC). The cell lines from ATCC were grown in either RPMI-1640 (Gibco) or DMEM (Gibco) with FBS, GlutaMax (Gibco), and penicillin/streptomycin at 37°C, 5% CO_2_. Tunicamycin (T7765) and 2-DG (25972) were purchased from Millipore-Sigma.

### Western blot

Antibodies against TFEB (4240), 4E-BP1 (9644), p-4E-BP1 (S65) (9451), p-4E-BP1 (T37/46) (2855), p-Rictor (2114), p-Rictor (T1135) (3806), p-ACC (3661), ACC (3676), AKT (2920), p-AKT (S473) (4060), LC3 (12741), p62 (SQSTM1) (88588), Ero1-Lα (3264), BiP (3177), IRE1α (3294), PDI (3501), CHOP (2895), PERK (5683), MYC (2276), GAPDH (5174), Histone H3 (14269), and Lamin A/C (4777) were purchased from Cell Signaling Technology. Total cell lysates were prepared using ice-cold RIPA buffer (Thermo Scientific) supplemented with Halt^TM^ Protease and Phosphatase Inhibitor Cocktail (Thermo Scientific). Protein concentration was quantified using the DC Protein Assay (Bio-Rad). Equal amounts of lysates were loaded onto the SDS-polyacrylamide gel and subsequently transferred to a PVDF/0.2μm nitrocellulose membrane (Bio-Rad). Blots were incubated overnight with primary antibodies diluted at 1:1000 in 5% non-fat milk at 4°C. Secondary antibodies anti-rabbit IgG-HRP (Cell Signaling Technology, 7074), StarBright Blue 700 anti-rabbit IgG (Bio-Rad, 12004162), StarBright Blue 700 anti-mouse IgG (Bio-Rad, 12004159), or DyLight 488 anti-mouse IgG (Bio-Rad, STAR117D488GA) were diluted in 5% non-fat milk. Either GAPDH (Cell Signaling Technology, 5174) or hFAB Rhodamine anti-Tubulin (Bio-Rad, 12004165) was used as a loading control.

### Plasmids and transfection

The following plasmids were obtained from Addgene: psPAX2 (12260; Dr. Didier Trono) and pCMV-VSV-G (8454; Dr. Bob Weinberg). TFEB (TRCN0000013109; TRCN0000013108) and HSPA5 (TRCN0000001024) shRNAs were purchased from Millipore-Sigma. TFEB (S142A) was a gift from Dr. Andrea Ballabio at Telethon Institute of Genetics and Medicine, Italy. TFEB (S142A) was cloned into pLenti-C-Myc-DDK-IRES-Puro Lentiviral Gene Expression Vector (OriGene). HEK293T cells were transfected using Lipofectamine 3000 (Invitrogen), and lentiviral particles were harvested to transduce target cell lines.

### Microscopy and immunocytochemistry

Brightfield (BF) imaging was done on Olympus CKX41 inverted microscope. Paraffin-embedded samples were sectioned, stained with Hematoxylin and Eosin (H&E), and imaged on Nikon Eclipse Ni-E upright microscope.

Immunocytochemistry (ICC) was performed as previously described (Soleimani *et al*., 2022). Primary antibody LC3 (Cell Signaling Technology, 12741) was used in 5% goat serum at 4°C overnight. The secondary antibody Alexa Fluor 555 anti-rabbit IgG (Cell Signaling Technology, 4413) was used in 5% goat serum for 2h at room temperature. Next, cells were counter-stained with DAPI and imaged using Nikon Eclipse Ti-2 A1R confocal microscope.

### Flow cytometry

Cells were trypsinized and washed with the flow wash buffer comprising PBS supplemented with FBS and EDTA. Primary antibodies against CD44 (BioLegend, 338806), CD24 (BioLegend, 311104), and CD49f (BioLegend, 313612) were diluted in the flow wash buffer and incubated with cells for 30min at room temperature. After washing with the flow wash buffer, cells were resuspended in propidium iodide and analyzed by flow cytometry (Cytek Aurora).

### qRT-PCR

Total RNA was extracted from cells using the PureLink RNA Mini Kit (Invitrogen), and cDNA was synthesized using the iScript cDNA Synthesis Kit (Bio-Rad). qRT-PCR was performed using the iTaq Universal SYBR Green Supermix (Bio-Rad) on a CFX384 RT-PCR detection system (Bio-Rad).

### Mammosphere formation assay

The mammosphere culture medium was prepared with DMEM/F12 supplemented with GlutaMax, penicillin/streptomycin, EGF, bFGF, and B27. Cells were trypsinized and plated in 24-well ultra-low attachment plates for 7-10 days. Primary mammospheres were trypsinized and re-plated in 24-well ultra-low attachment plates to form secondary mammospheres.

### Clonogenic assay

Cells were washed, trypsinized, and counted. Either 6-well or 12-well plates were seeded with 2.5×10^3^ or 1.5×10^3^ cells/well, respectively. Plates were incubated for 7-10 days at 37°C, 5% CO_2_. Colonies were stained with crystal violet and imaged. Colonies were solubilized in 10% acetic acid to quantify clonogenic capacity and read for absorbance at 590nm in a plate reader (BioTek).

### Gene overexpression and shRNA knockdown

Cells were subjected to gene knockdown using shRNAs targeting TFEB or HSPA5 via lentiviral transduction. A scramble shRNA was used as a control in all shRNA-mediated knockdown experiments. Overexpression of TFEB (S142A) and RagC (S75L) was carried out via lentiviral transduction. An empty pLenti-C-Myc-DDK-IRES-Puro vector was used as a control in all overexpression experiments.

### Limiting dilution tumor initiation assay and tumor xenograft assay

Female NSG (NOD.Cg-PrkdcSCID Il2rgtm1Wjl/SzJ) mice, purchased from the Jackson Laboratory, were orthotopically injected with HCC1806 cells expressing either scramble control or TFEB shRNA. Each cell line was injected with serial dilutions 5×10^5^(n=5), 5×10^4^(n=6), 5×10^3^(n=7), or 5×10^2^(n=9). Tumor growth in mice was tracked for 140 days following injection. The Tumor Initiating Cell (TIC) frequency for each cell line was calculated using ELDA (Extreme Limiting Dilution Analysis) (Hu & Smyth, 2009) at https://bioinf.wehi.edu.au/software/elda.

Female athymic (Foxn1^nu/nu^) mice, purchased from Envigo, were orthotopically injected with 1×10^6^ cells transduced with either scramble control or TFEB shRNA. Tumors were measured three times a week using a caliper, and tumor volume was computed as follows: (length×width^2^/2).

### Statistical analysis

Statistical analyses were carried out using GraphPad Prism 9. The specific statistical tests used included student’s t-test, one-way ANOVA followed by Dunnett’s post-hoc, and two-way ANOVA followed by Sidak’s post-hoc as denoted.

### Data availability

The authors generated the data, which are available upon request.

## ACKNOWLEDGMENTS

We thank the DNA Sequencing Facility Core at UT Austin for Sanger sequencing, the Center for Biomedical Research Support at UT Austin for flow cytometry and microscopy, and all the Developmental Therapeutics Laboratory personnel for their guidance and support.

## The Paper Explained Problem

TNBC is an aggressive subtype of breast cancer that is difficult to treat due to its lack of effective molecular targets. Triple-negative breast CSCs are thought to be responsible for the recurrence of TNBC after initial treatment. Cancer cells rely on glycolysis to support their rapid growth. Cancer stem cells can adapt to metabolic and oxidative stress in the tumor microenvironment. It is not well understood how TNBC stem cells survive under conditions of metabolic stress.

## Results

In this study, we investigated the role of a transcription factor called TFEB in the survival of TNBC stem cells under 2-DG-induced stress. We found that TFEB expression was higher in TNBC than in other breast cancer subtypes. TFEB was necessary for TNBC mammosphere formation *in vitro* and tumor formation *in vivo*. 2-DG-induced stress suppressed self-renewal in TNBC, consistent with previous relevant studies. Overexpressing constitutively nuclear TFEB rescued mammosphere formation in 2-DG-treated cells. Additionally, we showed that TFEB mediates 2-DG-induced UPR and autophagy as potential protective mechanisms in response to 2-DG.

### Impact

In this study, we shed light on how TFEB protects TNBC stem cells under metabolic stress. The findings suggest that TFEB modulates major metabolic pathways in CSCs that may contribute to drug resistance and disease relapse. Overall, this study provides valuable insights into the regulation of triple-negative breast CSCs and their survival mechanisms in response to stress. Understanding these TFEB-regulated pathways could reveal potential therapeutic opportunities for TNBC.

## FIGURE LEGENDS

**Figure EV1.** Effect of 2-DG on TNBC *in vitro.* **A)** Western blot analysis of scramble or TFEB KD HCC1806, MDA-MB-231, and MDA-MB-157 cells. **B)** Mammosphere formation assay of MDA-MB-231 transduced with either scramble or TFEB shRNA and grown in mammosphere media for 7-10 days, followed by passaging to form secondary mammospheres. Scale bar: 200μm. **C)** Clonogenic assay of MDA-MB-231 transduced with either scramble or TFEB shRNA.

**Figure EV2.** 2-DG suppresses cancer stem cell phenotype in TNBC. **A)** Clonogenic assay of BT549 and HCC38 treated with either vehicle or 2-DG (1, 2, 5, 10 & 20mmol/L) for 72h and recovered for 7-10 days. **B)** Flow cytometric analysis of CD44^high^/CD24^low^ in HCC1937 treated with either vehicle or 2-DG for 24h. (one-way ANOVA, Dunnett test) *, *p* < 0.05; **, *p* < 0.01; ***, *p* < 0.001; ****, *p* < 0.0001

**Figure EV3.** 2-DG induced TFEB nuclear translocation and activates AMPK signaling. **A**) Western blot analysis of nuclear/cytosolic fractions of HCC1806 treated with either vehicle or 2-DG for 24h. Tubulin was used a cytosolic marker, and Lamin A/C as a nuclear marker. **B)** Western blot analysis of indicated cell lines glucose starved for 3h, 8h, or 24h. **C**) Western blot analysis of TFEB, p-4E-BP1 (S65), p-4-EBP1 (T37/46), p-p70S6K (T389), p-Rictor (T1135), p-AKT (S473), and p-ACC (S79) in indicated cell lines treated with either vehicle or 2-DG for 24h. **D)** Scatter plots of TFEB vs EIF2AK, TFEB vs DDIT3, and TFEB vs HSPA5 mRNA expression in breast cancer tumors as retrieved by Correlation AnalyzeR. Veh: Vehicle; Glc: Glucose

